# CD-HIT-OTU-MiSeq, an Improved Approach for Clustering and Analyzing Paired End MiSeq 16S rRNA Sequences

**DOI:** 10.1101/153783

**Authors:** Weizhong Li, Yuanyuan Chang

**Affiliations:** J Craig Venter Institute, La Jolla, California, United States of America

## Abstract

In recent years, Illumina MiSeq sequencers replaced pyrosequencing platforms and became dominant in 16S rRNA sequencing. One unique feature of MiSeq technology, compared with Pyrosequencing, is the Paired End (PE) reads, with each read can be sequenced to 250-300 bases to cover multiple variable regions on the 16S rRNA gene. However, the PE reads need to be assembled into a single contig at the beginning of the analysis. Although there are many methods capable of assembling PE reads into contigs, a big portion of PE reads can not be accurately assembled because the poor quality at the 3’ ends of both PE reads in the overlapping region. This causes that many sequences are discarded in the analysis. In this study, we developed a novel approach for clustering and annotation MiSeq-based 16S sequence data, CD-HIT-OTU-MiSeq. This new approach has four distinct novel features. (1) The package can clustering PE reads without joining them into contigs. (2) Users can choose a high quality portion of the PE reads for analysis (e.g. first 200 / 150 bases from forward / reverse reads), according to base quality profile. (3) We implemented a tool that can splice out the target region (e.g. V3-V4) from a full-length 16S reference database into the PE sequences. CD-HIT-OTU-MiSeq can cluster the spliced PE reference database together with samples, so we can derive Operational Taxonomic Units (OTUs) and annotate these OTUs concurrently. (4) Chimeric sequences are effectively identified through *de novo* approach. The package offers high speed and high accuracy. The software package is freely available as open source package and is distributed along with CD-HIT from http://cd-hit.org. Within the CD-HIT package, CD-HIT-OTU-MiSeq is within the usecase folder.

## 1 Introduction

One of the fundamental questions in microbiome studies is to estimate the microbial diversity in the environment. And the most common approach is to measure the 16S rRNA genes in the samples using amplicon sequencing approach established and developed during the last decade. In the earlier studies, Pyrosequencing (i.e. the 454 sequencing) is the major sequencing platform, which underwent several generations of platform with each newer platform offering longer reads. After the discontinuation of the 454 sequencing platforms in 2013, Illumina’s MiSeq became the dominant platform for 16S rRNA amplicon sequencing.

The 16S rRNA sequence data are usually analyzed by clustering-based approach to derive Operational Taxonomic Units (OTUs), which describe the distinct groups of microbial organisms at different taxonomic level. OTUs clustered at 97% sequence identity are usually used by the field to represent distinct species. However, noise in PCR-based amplification, sequencing errors and artifacts often cause overestimation of OTUs [1, 2]. So, in the past, many methods and protocols were developed to identify these errors and to reduce the false OTUs. For the data from Pyrosequencing platforms, the most adopted methods was through flowgram clustering and denoising, which were implemented in programs such as PyroNoise [3], Denoiser [4] and AmpliconNoise [5]. In addition, most methods used strict quality filtering and trimming on the raw sequencing reads and some methods also deployed pre-clustering process, as introduced in SLP [1]. In order to identify chimeric reads, both reference-based methods such as ChimeraSlayer [6] and *de novo* approaches (e.g. UCHIME [7]) were introduced. For sequences from MiSeq platform, although the flowgram based denoising methods are no longer applicable, but many other techniques and protocols developed for 454 data, such as strict quality filtering and pre-clustering are still applied in MiSeq data analysis. These methods are all available from many commonly used 16S pipelines, such as Mothur [8] and Qiime [9].

One unique feature of MiSeq technology, compared to Pyrosequencing, is the Paired End (PE) reads, with forward (R1) and reverse (R2) read can be sequenced to 250-300 bases. Therefore, the whole PE reads can cover multiple variable regions on the 16S rRNA gene. However, the PE reads need to be assembled into a single contig at the beginning of the analysis. MiSeq, like some other Illumina’s sequencers, produces relative lower quality bases towards the end of the reads. Also R2 reads usually have more errors than R1 reads. So many PE reads cannot be assembled perfectly without mismatching base. So, methods have been developed to join the PE reads to produce higher quality contigs permitting erroneous and mismatching bases. PANDAseq [10] and PEAR [11] are two of such programs that can effectively assemble a large number of PE reads. Other programs that can join the PE reads include FLASH [12] and COPE [13]. In addition, pipelines such as Mothur and Qiime all have built-in tool to assemble PE reads into contigs. Despite there are many tools for PE read assembly, for some datasets, a big portion of reads can not be assembled because the poor quality at the 3’ end of both PE reads in the overlapping region. Even if the contigs can be assembled allowing many mismatches in the overlapping region, these contigs may have too many errors to be used. In fact, discarding low quality contigs are standard step in programs like Mothur.

In the past, we developed ultra-fast sequence clustering tool CD-HIT [14-17], which were used to cluster 16S sequences in many applications. In order to address the problem of overestimation of OTUs due to sequence errors in Pyrosequencing data, we developed CD-HIT-OTU pipeline [18], with high speed and accuracy. Here we present another novel approach that based on CD-HIT package for clustering and annotating MiSeq based 16S sequence data, CD-HIT-OTU-MiSeq. This new approach has four distinct novel features. (1) The recently released CD-HIT package can cluster PE reads without the requirement for joining PE reads into contigs, so the CD-HIT-OTU-MiSeq can work with PE reads that can not be effectively assembled. (2) A user can select and analyze only high quality portion of the PE reads, such as first 200 base from R1 reads and first 150 base from R2 reads, according to sequencing base quality profile. (3) We implemented a tool that can splice out the target region (e.g. V3-V4) from a full-length 16S rRNA reference sequence database into the PE sequences. CD-HIT-OTU-MiSeq can cluster the spliced PE reference database together with the sample, so we can derive OTUs and annotate these OTUs concurrently. (4) Chimeric sequences are effectively identified through *de novo* approach. In addition, CD-HIT-OTU-MiSeq adopted other denoising approaches from our earlier CD-HIT-OTU.

Our approach provides an alternative way for analyzing MiSeq 16S data, especially the datasets where a considerable portion cannot be assembled into contigs. The software package is freely available and is distributed along with CD-HIT package from http://cd-hit.org. Within the CD-HIT package, CD-HIT-OTU-MiSeq is within the usecase folder.

## 2 Methods

### 2.1 Overall Clustering Process

CD-HIT-OTU-MiSeq includes three major processes, reference database preparation, sequence quality control (QC), and OTU clustering and annotation.

The most important unique feature of this method is to only use high quality region at the 5’ ends of R1 and R2 reads. For example, the effective clustering read length can be 200 bases for R1 and 150 bases for R2. The effective portions of PE reads are clustered together with spliced PE sequences from the reference database to derive OTUs (Figure 1). In this paper, we will show the results of OTUs based on different effective clustering read lengths. In practice, programs such as FASTQC (http://www.bioinformatics.babraham.ac.uk/projects/fastqc/) can be used to scan the raw reads to help choose the effective clustering read length of R1 and R2.

**Figure 1.**
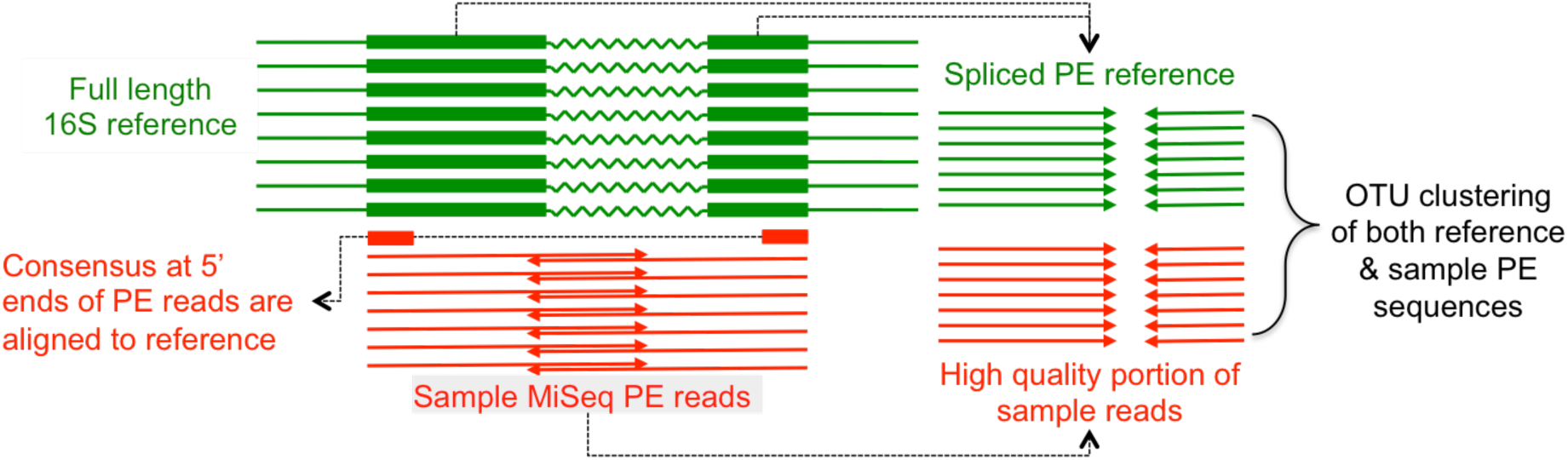
Clustering of high quality portion of PE reads together with spliced PE sequences from 16S reference database

### 2.2 Reference database preparation

We implemented a tool that can splice out the target amplicon region (e.g. V3-V4) from a full-length 16S rRNA reference sequence database, such as Greengene [19], RDP [20] and Silva [21], into PE sequences. Given a Miseq PE dataset, this tool performs the following steps.

1. It scans the 5′ ends of both R1 and R2 reads to get consensus sequences of at least 30 bases.
2. Cd-hit-est-2d (parameters: -c 0.8 -n 5 -r 1 -p 1 -b 5 -G 0 -A 30 -s2 0.01) is used to align the consensus sequence to the full-length 16S reference sequences.
3. Two fragments from each full-length 16S reference sequence were cut out at the aligned position. The size of forward and reverse fragments can be selected to match the effective clustering read length. The reverse fragments are converted into complementary sequences and both fragments are saved in PE fasta files, which are compatible with MiSeq PE sequence files.
4. This spliced PE sequences are clustered at 99% identity to remove redundant sequences with cd-hit-est (parameters: -c 0.99 -n 10 -p 1 -b 5 -G 1 -g 1 -P 1 -l 11 -sc 1).

If there are multiple samples in a project sequenced with the same amplicon of same variable region, only one spliced reference database is needed.

### 2.3 QC

In this study, the raw reads are processed with Trimmomatic [22] to trim low-quality bases and to filter out low quality reads (parameters: SLIDINGWINDOW:4:15 LEADING:3 TRAILING:3 MINLEN:100 MAXINFO:80:0.5). With this setting, Trimmomatic uses a 4-base sliding window to trim a read at window position where average quality score is below 15. The PE reads are kept if both are at least 100 bases after trimming.

### 2.4 OTU clustering and annotation

Although only the high quality portion of (up to user-selected effective clustering read lengths) PE reads are used in clustering. There is no need to physically trim the PE reads to effective clustering read length since cd-hit-est program has option to use only the user-selected portion of sequences. OTU clustering has following steps:

1. PE reads are clustered with cd-hit-est at 100% identity to find clusters of exact duplicated PE reads (parameters: -sf 1 -sc 1 -P 1 -r 0 -cx effective_read_length_R1 -cy effective_read_length_R2 -c 1.0 -n 10 -G 1 -b 1 -d 0 -p 1). The resulting unique PE reads are saved in decreasing order of abundance (enabled by parameter -sf 1). Clusters with greater number of exact duplicates are much more likely to be sequencing error free.
2. Unique PE reads from step 1 are clustered at 99% identity (cd-hit-est parameters: -P 1 -r 0 -cx effective_read_length_R1 -cy effective_read_length_R2 -c 0.99 -n 10 -G 1 -b 1 -d 0 -p 1). Since the input PE reads are in decreasing order of abundance, during the cd-hit-est clustering process, the most abundant unique PE reads (also the most likely error free reads) will form clusters to recruit less abundant PE reads with <=1% errors.
3. R1 reads only from the unique PE reads are clustered at 99% identity (cd-hit-est parameters: -c 0.99 -n 10 -cx 75 -G 0 -b 1 -d 0 -p 1 -A 50). R2 reads are also separately clustered with same parameters. Similar to step 2, less abundant unique reads will be clustered into more abundant unique reads.
4. Using the clustering results from step 1-3, chimeric sequences are identified using the same way as we previously implemented in cd-hit-dup [18]. PE reads that are clustered to another PE reads in step 2 are not chimeric. For a remaining PE reads C, if C.R1 and C.R2 are clustered into A.R1 and B.R2 and A and B are not paired and both A and B’s abundance is more than twice of C’s abundance, then C is considered chimeric. This procedure is also similar to Uchime [7].
5. Given an abundance cutoff (e.g. 0.0001), the small clusters with fewer than N sequences are filtered out. Here N = cutoff * number of high quality PE reads.
6. The representative sequences from step 2, excluding chimeric reads identified in step 4 and small clusters found in step 5 are clustered at 97% identity (cd-hit-est parameters: -P 1 -r 0 -cx effective_read_length_R1 -cy effective_read_length_R2 -c 0.97 -n 10 -G 1 -b 10 -d 0 -p 1). The generated clusters are OTUs.
7. Cd-hit-est-2d is used to recruit spliced reference to OTU clusters generated in step 6 (cd-hit-est-2d parameters: -P 1 -r 0 -cx effective_read_length_R1 -cy effective_read_length_R2 -c 0.97 -n 10 -G 1 -b 10 -d 0 -p 1). There may be multiple reference sequences in the a single OTU cluster, only the sequence most similar to the representative sequence in that OTU is kept, and is used to annotate the OTU.

### 2.5 OTU analysis of multiple Miseq samples

In most experiments, multiple samples were studied using the same protocol and same amplicon. It is effective to pool the samples together and cluster them to a derive OTU table that are comparable across samples.

For multiple samples, only one spliced reference database is needed. Each sample can be processed individually for QC and OTU clustering. Then the non-chimeric non-small clusters of all samples are pooled and are clustered at 97% identity (same as step 6 in 2.4) and the spliced reference database are recruited (same as step 7 in 2.4).

## 3 Results

### 3.1 Mock datasets

In this study, we used several Mock datasets sequenced with MiSeq platform. These Mock samples have been used to validate tools and methods analyzing 16S sequences. The First Mock community (Mock 1) is composed of 21 bacterial isolates, available from BEI Resources (HM-278D v3.1). Sequence data for Mock 1 were from study [23], which sequenced V4, V34 and V45 regions in multiple runs. Data from 4 runs (130401, 130403, 130417, 130422) were used in this study. The second Mock community (Mock 2) is an earlier version of Mock1 (BEI Resources, HM-276D, Genomic DNA from Microbial Mock Community B, even concentration). Mock 2 contains 20 bacteria strains. For Mock 2, there were two runs for V4 and one run for V45 [24]. The third Mock community (Mock 3) contains 12 species. Three runs were performed on V34 region [25]. So total 18 Mock samples were used in this study (Table 1). Mock 1 datasets were downloaded from https://www.mothur.org/MiSeqDevelopmentData.html. Mock 2 datasets were downloaded from EMBL-EBI ENA under accession PRJEB4688 and Mock 3 datasets were downloaded from NCBI SRA under accession SRP066114. Table 1 also shows the number of PE reads and number of high quality PE reads after QC.

**Table 1.**
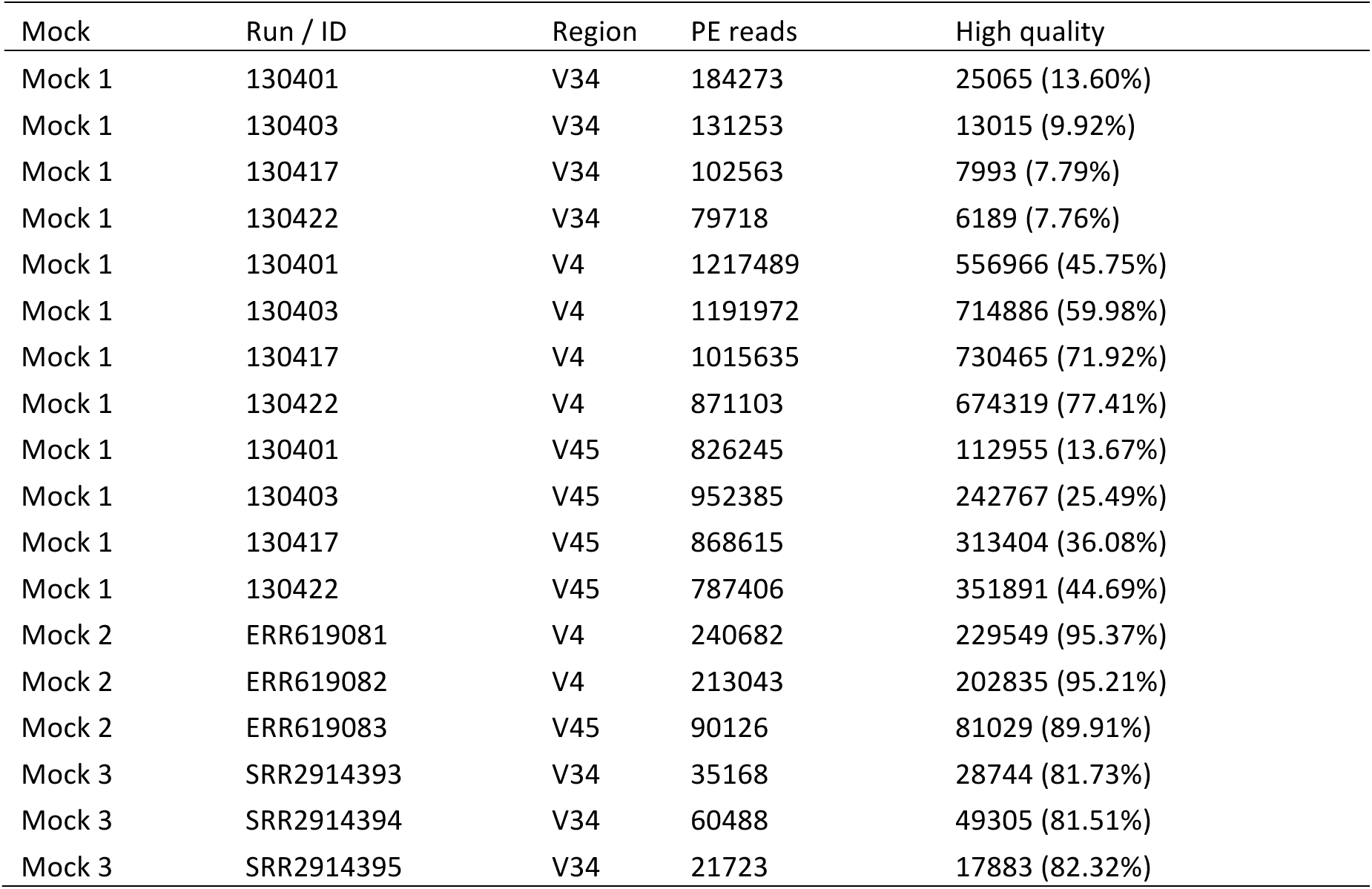
Mock datasets used in this study

| Mock | Run / ID | Region | PE reads | High quality |
| --- | --- | --- | --- | --- |
| Mock 1 | 130401 | V34 | 184273 | 25065 (13.60%) |
| Mock 1 | 130403 | V34 | 131253 | 13015 (9.92%) |
| Mock 1 | 130417 | V34 | 102563 | 7993 (7.79%) |
| Mock 1 | 130422 | V34 | 79718 | 6189 (7.76%) |
| Mock 1 | 130401 | V4 | 1217489 | 556966 (45.75%) |
| Mock 1 | 130403 | V4 | 1191972 | 714886 (59.98%) |
| Mock 1 | 130417 | V4 | 1015635 | 730465 (71.92%) |
| Mock 1 | 130422 | V4 | 871103 | 674319 (77.41%) |
| Mock 1 | 130401 | V45 | 826245 | 112955 (13.67%) |
| Mock 1 | 130403 | V45 | 952385 | 242767 (25.49%) |
| Mock 1 | 130417 | V45 | 868615 | 313404 (36.08%) |
| Mock 1 | 130422 | V45 | 787406 | 351891 (44.69%) |
| Mock 2 | ERR619081 | V4 | 240682 | 229549 (95.37%) |
| Mock 2 | ERR619082 | V4 | 213043 | 202835 (95.21%) |
| Mock 2 | ERR619083 | V45 | 90126 | 81029 (89.91%) |
| Mock 3 | SRR2914393 | V34 | 35168 | 28744 (81.73%) |
| Mock 3 | SRR2914394 | V34 | 60488 | 49305 (81.51%) |
| Mock 3 | SRR2914395 | V34 | 21723 | 17883 (82.32%) |

### 3.2 OTU clustering

Following the procedures described in the method, the 18 Mock samples were clustered at 97% identity to derive OTUs at species level. In this analysis, Greengene was used as reference database. Five pairs of effective clustering read lengths (225, 175), (200, 150), (175, 125), (150, 100) and (125, 75) were selected for samples sequenced at V34 or V45. Two pairs of effective clustering read lengths (150, 100) and (125, 75) were used for samples of V4 region. Two abundance cutoffs were used: 0.0001 and 0.0005. The numbers of OTUs are shown in Table 2.

**Table 2.** OTUs calculated at different effective clustering read lengths and different abundance cutoffs

| Run / ID | Region | OTUs<br>225, 175 | OTUs<br>200,150 | OTUs<br>175, 125 | OTUs<br>150, 100 | OTUs<br>125, 75 |
| --- | --- | --- | --- | --- | --- | --- |
| Mock 1 |  |  |  |  |  |  |
| 130401 | V34 | 30,20 | 35,20 | 29,19 | 29,19 | 24,19 |
| 130403 | V34 | 38,19 | 38,19 | 37,20 | 45,21 | 41,22 |
| 130417 | V34 | 36,21 | 38,21 | 36,21 | 33,22 | 33,22 |
| 130422 | V34 | 50,26 | 50,27 | 52,31 | 56,33 | 60,36 |
| 130401 | V4 |  |  |  | 24,20 | 24,20 |
| 130403 | V4 |  |  |  | 25,20 | 27,20 |
| 130417 | V4 |  |  |  | 24,20 | 26,20 |
| 130422 | V4 |  |  |  | 27,20 | 28,20 |
| 130401 | V45 | 30,21 | 27,20 | 25,20 | 29,20 | 30,20 |
| 130403 | V45 | 21,21 | 21,21 | 21,21 | 25,21 | 29,21 |
| 130417 | V45 | 23,21 | 24,21 | 24,21 | 27,21 | 30,21 |
| 130422 | V45 | 21,21 | 22,21 | 22,21 | 24,21 | 25,21 |
| Mock 2 |  |  |  |  |  |  |
| ERR619081 | V4 |  |  |  | 55,23 | 50,22 |
| ERR619082 | V4 |  |  |  | 40,22 | 40,20 |
| ERR619083 | V45 | 40,18 | 33,21 | 39,23 | 32,18 | 42,19 |
| Mock 3 |  |  |  |  |  |  |
| SRR2914393 | V34 | 29,12 | 22,12 | 26,13 | 33,14 | 28,14 |
| SRR2914394 | V34 | 19,12 | 23,12 | 25,12 | 33,12 | 26,12 |
| SRR2914395 | V34 | 29,12 | 27,12 | 28,12 | 35,12 | 32,12 |

Each cell of this table shows number of OTUs at abundance cutoffs 0.0001 and 0.0005.

At different effective clustering read lengths, the number of OTUs slightly or moderately fluctuates, but is not correlated with the effective clustering read length either positively or negatively. The number of OTUs returned by our method is very close to the truth: 21, 20 and 12 for Mock 1, 2 and 3, especially at cutoff 0.0005. Mock 1 samples were originally studied in [23], the average number OTUs derived from our study based on the whole data sets (up to 1.2 million) at cutoff 0.0001 and 0.0005 are 31 and 22 respectively, comparable to or less than the OTUs calculated in reference [23], which range from 22 to 192, based on rarefaction of each sample to 5,000 sequences per sample (Table 2 of reference [23]).

Mock 2 were initially analyzed in reference [24]; the reported OTUs for the 3 samples using Qiime are from 138 to 143 (Table 2 in reference [24]), which are much more than the OTUs identified by our method. Mock 3 samples were first analyzed in previous study [25], where the reported OTUs by several different methods range from 50 to 148 (Table S12 in reference [25]), which are higher than our results.

In fact, study [25] analyzed both Mock 1, 2 and 3 samples and reported the number of OTUs in Table S12 using several methods including USEARCH, UNoise, Mothur and IPED. Compared to that, our OTUs are constantly lower than the reported results, except that USEARCH and UNoise performed better on Mock 1 V34 samples.

So, compared to the published results from multiple studies [23] [24][25] with many different methods, the OTUs by our method are generally more accurate.

### 3.3 Compute time

The compute time for clustering these Mock samples varied from a few to ~10 minutes per sample, depending on the sample size and effective clustering read length considered. Our approach is much faster than other popular methods including Mothur and Qiime. Because of the ultra-high speed, CD-HIT-OTU-MiSeq is able to process a hundred sample of similar size in a couple of hours.

### 3.4 OTU Annotation

In our process, sequences were annotated if they were clustered together with the reference 16S genes. We checked the clusters, in all cases, the known bacteria species included in the Mock 1 and Mock 2 samples were found in the OTUs. It is expected, in a few cases, very closely related species are clustered in to the same OTUs, with effective clustering read length of (125, 75). For Mock 3, we have difficult to find the species composition from the reference [25], but it is very clear that top large OTUs are corresponding to these species that constitute the Mock 3 samples.

Besides the large OTUs with known species, the remaining small OTUs are either cluster of sequences with larger sequencing errors, or from contaminating microbes at very low abundance. We observed both sequences for all Mock samples.

## 4 Conclusion

CD-HIT-OTU-MiSeq gives an alternative way for analyzing MiSeq 16S data, without assembling the PE reads into contigs. It is especially useful when a considerable portion of PE reads cannot be assembled into contigs without mismatch. With further improvements from our previous CD-HIT-OTU, this package offers high accuracy and speed in OTU clustering and is able to process hundreds of MiSeq samples in hours.

The software package is freely available and is distributed along with CD-HIT package from http://cd-hit.org. Within the CD-HIT package, CD-HIT-OTU-MiSeq is within the usecase folder. The detailed document and users’ guide are available from the package.

